# White Matter Tract Vulnerability to Amyloid Pathology on the Alzheimer’s Disease Continuum

**DOI:** 10.1101/2025.08.18.670970

**Authors:** Bramsh Qamar Chandio, Talia M. Nir, Julio E. Villalon-Reina, Sophia I. Thomopoulos, Yixue Feng, Robert I. Reid, Clifford R. Jack, Michael W. Weiner, Eleftherios Garyfallidis, Neda Jahanshad, Meredith N. Braskie, Sid O’Bryant, Paul M. Thompson, the Alzheimer’s Disease Neuroimaging Initiative, the Health and Aging Brain Study (HABS-HD) Study Team

## Abstract

Alzheimer’s disease (AD) is marked by progressive cognitive decline and memory loss, due to the abnormal accumulation of amyloid-beta (*Aβ*) plaques, which facilitate the spread of tau pathology, and a gradually spreading pattern of neuronal loss. Understanding how amyloid positivity affects the brain’s neural pathways is critical for understanding how the brain changes with AD pathology. Tractometry offers a powerful approach for the *in vivo*, 3D quantitative assessment of white matter tracts, enabling the localization of microstructural abnormalities in diseased populations and those at risk. In this study, we applied BUAN (Bundle Analytics) tractometry to multi-cohort diffusion MRI data from a total of 1,908 participants: 606 participants in ADNI3 (Alzheimer’s Disease Neuroimaging Initiative Phase 3) and 1,302 participants from the HABS-HD (Health and Aging Brain Study–Health Disparities). Using BUAN and along-tract statistical analysis, we assessed the localized effects of amyloid positivity on white-matter pathways, which may be further influenced by downstream tau accumulation. Amyloid positivity was quantified via amyloid-sensitive positron emission tomography (PET). BUAN enables tract-specific quantification of white matter microstructure and supports statistical testing along the full length of fiber bundles to detect subtle, spatially localized associations. We present 3D visualizations of tractwise amyloid associations, highlighting distinct patterns of white matter degeneration in AD.

## I. Introduction

Alzheimer’s disease (AD) is characterized by progressive cognitive decline, with hallmark pathologies including amyloid-beta (*Aβ*) plaque accumulation and tau neurofibrillary tangles [1]–[3]. While these abnormalities are typically observed in gray matter, growing evidence shows that white matter (WM) microstructure is also disrupted in AD, potentially reflecting early or propagating disease processes [4], [5]. In particular, *Aβ* pathology is associated with myelin degradation, neuroinflammation, and axonal injury, contributing to widespread disconnection across the brain’s networks.

Diffusion MRI (dMRI) enables non-invasive mapping of WM microstructure by capturing the directional movement of water molecules in tissue [6]. Tractography reconstructs white matter pathways, and tractometry further quantifies microstructural properties along these pathways, offering a high-resolution approach to detect disease effects [7], [8]. While voxel-wise techniques such as tract-based spatial statistics (TBSS) have been used to relate WM changes to AD biomarkers [9], [10], they lack the spatial precision offered by tractometry approaches.

In this study, we used BUAN (Bundle Analytics) [7], an advanced tractometry pipeline, to investigate the effects of amyloid, as an upstream factor that may facilitate subsequent tau-related degeneration, on white matter microstructure in a large, diverse sample from the Alzheimer’s Disease Neuroimaging Initiative Phase 3 (ADNI3) [10] and the Health and Aging Brain Study–Health Disparities (HABS-HD) [11] cohorts. BUAN maps dMRI derived microstructural metrics along the trajectory of major white matter tracts, enabling localized detection of microstructural alterations associated with *Aβ* accumulation. In this study, we use *Aβ* status to examine how microstructural changes vary along the length of these tracts. This work extends our prior research that identified tract-specific degeneration associated with amyloid positivity using ADNI3 data [12]. A major goal of the study was to understand whether effects strengthen as more data is added, including data from a more ancestrally diverse dataset than ADNI, and multiple dMRI scanning protocols.

## II. Methods

### A. Data Preprocessing

We analyzed multi-cohort diffusion MRI data from 1,908 participants: 606 from ADNI3 (age 55–95; 287M/319F; 173 MCI, 59 AD, 374 CN; 277 *Aβ*+, 329 *Aβ*–) acquired using 7 protocols (GE36, GE54, P33, P36, S127, S31, S55), and 1,302 from HABS-HD (age 49–91; 471M/831F; 348 MCI, 93 AD, 861 CN; 239 *Aβ*+, 1,062 *Aβ*–) scanned with a single protocol (SE204) on Siemens Skyra and Vida. Please see Table 1 and Table 2 for details on demographics and dMRI acquisition protocols.

**Table 1.**
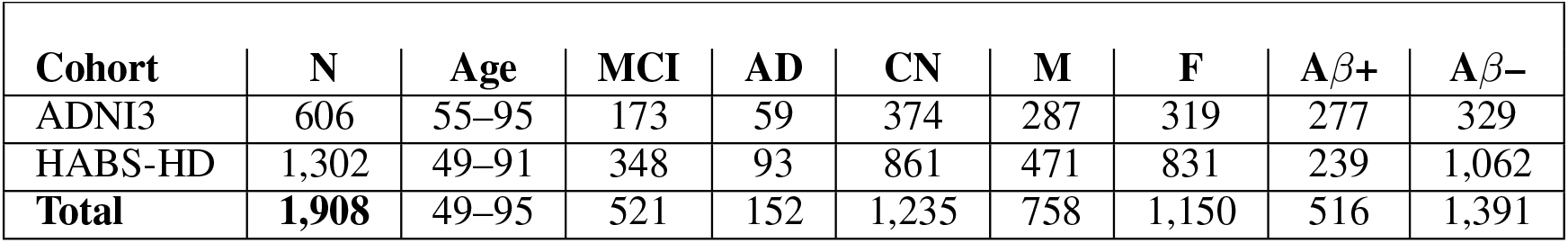
Participant Demographics.

**Table 2.**
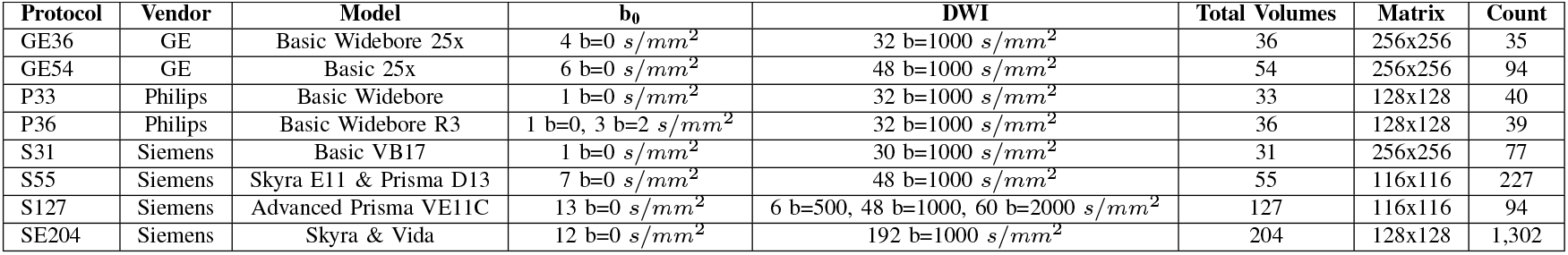
Diffusion MRI Acquisition Protocol Details.

| Protocol | Vendor | Model | b<sub>0</sub> | DWI | Total Volumes | Matrix | Count |
| --- | --- | --- | --- | --- | --- | --- | --- |
| GE36 | GE | Basic Widebore 25x | 4 b=0 s/mm<sup>2</sup> | 32 b=1000 s/mm<sup>2</sup> | 36 | 256x256 | 35 |
| GE54 | GE | Basic 25x | 6 b=0 s/mm<sup>2</sup> | 48 b=1000 s/mm<sup>2</sup> | 54 | 256x256 | 94 |
| P33 | Philips | Basic Widebore | 1 b=0 s/mm<sup>2</sup> | 32 b=1000 s/mm<sup>2</sup> | 33 | 128x128 | 40 |
| P36 | Philips | Basic Widebore R3 | 1 b=0, 3 b=2 s/mm<sup>2</sup> | 32 b=1000 s/mm<sup>2</sup> | 36 | 128x128 | 39 |
| S31 | Siemens | Basic VB17 | 1 b=0 s/mm<sup>2</sup> | 30 b=1000 s/mm<sup>2</sup> | 31 | 256x256 | 77 |
| S55 | Siemens | Skyra E11 & Prisma D13 | 7 b=0 s/mm<sup>2</sup> | 48 b=1000 s/mm<sup>2</sup> | 55 | 116x116 | 227 |
| S127 | Siemens | Advanced Prisma VE11C | 13 b=0 s/mm<sup>2</sup> | 6 b=500, 48 b=1000, 60 b=2000 s/mm<sup>2</sup> | 127 | 116x116 | 94 |
| SE204 | Siemens | Skyra & Vida | 12 b=0 s/mm<sup>2</sup> | 192 b=1000 s/mm<sup>2</sup> | 204 | 128x128 | 1,302 |

**Table 3.**
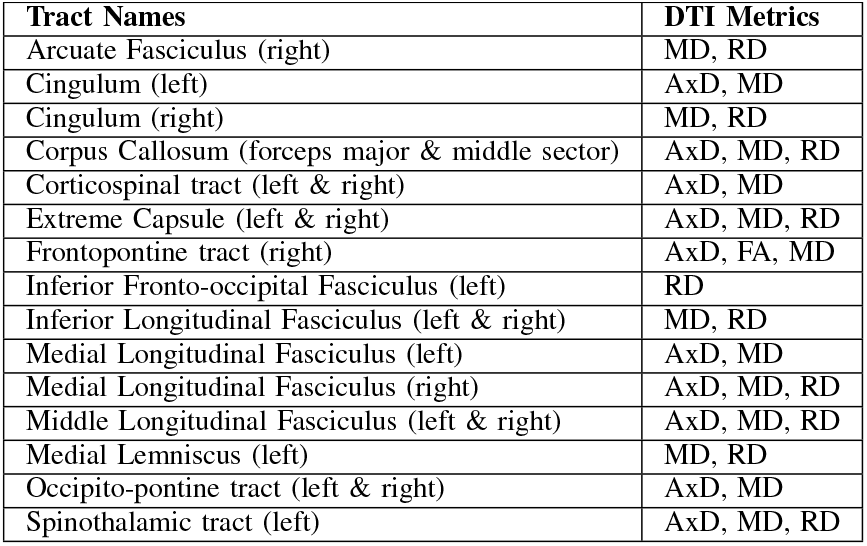
Tracts with Significant Amyloid Associations.

*Aβ* positron emission tomography (PET) status, i.e., positive or negative, was determined for each respective study. In ADNI, mean cortical standardized uptake value ratio (SUVR) from either 18F-florbetapir (*Aβ*+ defined as *>*=1.11) [13], [14] or florbetaben (*Aβ*+ defined as *>*=1.08) [15], [16] was used, with uptake normalized to the whole cerebellum reference region. In the HABS-HD, *Aβ*+ was defined as florbetaben SUVR *>*=1.08, consistent with the ADNI3 protocol.

Preprocessing of raw dMRI data involved several steps: denoising raw dMRI data using principal component analysis (PCA) for GE data, and Marchenko-Pastur PCA for Siemens and Philips data [17], [18]. Gibbs artifacts correction [19], and skull stripping [20]. Eddy currents and motion were corrected with additional corrections for slice-to-volume and outlier detection [21] and bias field inhomogeneities correction *dwibiascorrection*. The diffusion tensor imaging (DTI) model was used to extract 4 microstructural measures: fractional anisotropy (FA), mean diffusivity (MD), axial diffusivity (AxD), and radial diffusivity (RD).

### B. BUAN Tractometry

Fig. 1 illustrates the steps of the BUAN tractometry pipeline, along with visualizations of the process. We used Constrained Spherical Deconvolution (CSD) [22] for single-shell data and multi-shell multi-tissue CSD [23] for multishell data. Whole-brain tractograms were generated using probabilistic tractography. For tractography, seeds were placed in regions where FA *>* 0.15, with a seed density of 2 seeds per voxel and a step size of 0.5. We extracted 38 white matter (WM) tracts from the whole-brain tractograms using auto-calibrated RecoBundles [7], [24], leveraging model bundles from the HCP-842 tractography atlas [25].

**Fig. 1:**
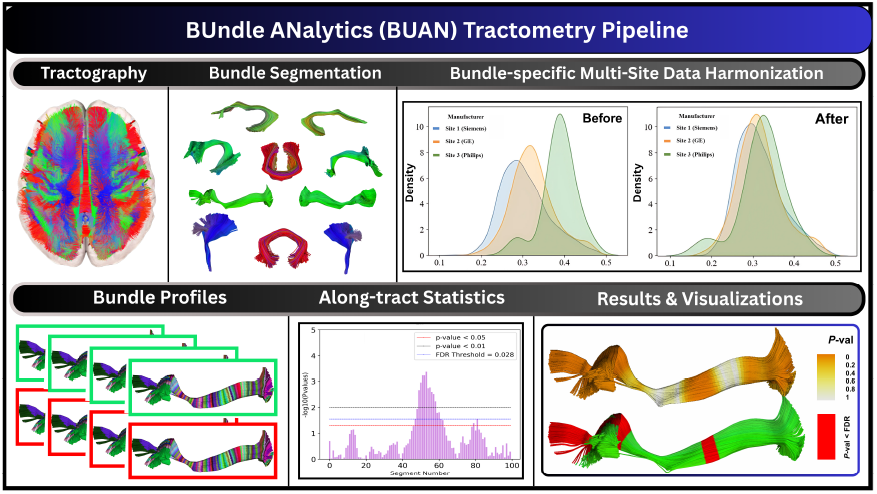
BUAN Tractometry Pipeline. The brain’s major neural pathways are reconstructed using diffusion MRI and tractography. Specific white matter tracts are extracted, and microstructural measures are projected onto them. Tract-specific data harmonization is applied to account for scanner/site-related variability. Bundle profiles are then statistically compared, enabling precise, localized analysis of white matter microstructure along brain pathways.

After extracting WM bundles, BUAN creates the bundle profiles for each bundle using 4 DTI-based microstructural metrics: FA, MD, RD, and AxD calculated in the diffusion native space. Bundle profiles are created by dividing the bundles into 100 horizontal segments using the model bundle centroids along the length of the tracts in common space. We clustered our model bundles using the QuickBundles [26] method to obtain a cluster centroid consisting of 100 points per centroid. We calculated Euclidean distances between every point on every streamline of the bundle and 100 points in the model bundle centroid. A segment number is assigned to each point in a bundle based on the shortest distance to the nearest model centroid point. Since the assignment of segment numbers is performed in the common space, we established the segment correspondence among subjects from different groups and populations. Microstructural measures such as FA are then projected onto the points of the bundles in native space.

Bundle profiles are harmonized using the ComBat method [27], [28] to correct for protocol/scanner effects as described in the harmonized BUAN tractometry pipeline [29]. We used Linear Mixed Models (LMMs) to test the effects of amyloid positivity on 38 white matter tracts. In each experiment, age and sex were modeled as fixed effects, and the protocol and subject were modeled as random terms. Although we harmonized the profiles with ComBat, we further accounted for protocol/scanner effects by modeling protocol as a random term in the LMM. Multiple testing correction was performed using the False Discovery Rate (FDR) [30] method at corrected *P*-value *<* 0.05. We perform tract-specific FDR correction for 100 tests per bundle.

## III. Results

Significant differences between *Aβ*+ and *Aβ*-groups were observed in the following white matter tracts (only tracts and DTI metrics that survived FDR correction are reported).

Across these tracts, amyloid positivity, which may be further influenced by downstream tau accumulation, was associated with increased diffusivity metrics (AxD, MD, RD) and decreased FA, indicating microstructural degeneration patterns consistent with established markers of Alzheimer’s pathology. Fig. 2 shows detailed *P*-value maps of the cingulum left (top) and cingulum right (bottom) bundles. In the left cingulum, stronger effects were detected using MD and AxD, while in the right cingulum, the effects were more prominent with MD and RD. The 3D tract visualizations in the first column of panels highlight regions with significant amyloid-related effects in red. *P*-value plots display the negative logarithm of *P*-value (y-axis) across 100 segments along each tract (x-axis). Cohort QQ-plots illustrate increased sensitivity to the statistical association when combining data from ADNI3 and HABS-HD, while protocol QQ-plots show enhanced detection power with the inclusion of multiple acquisition protocols.

**Fig. 2:**
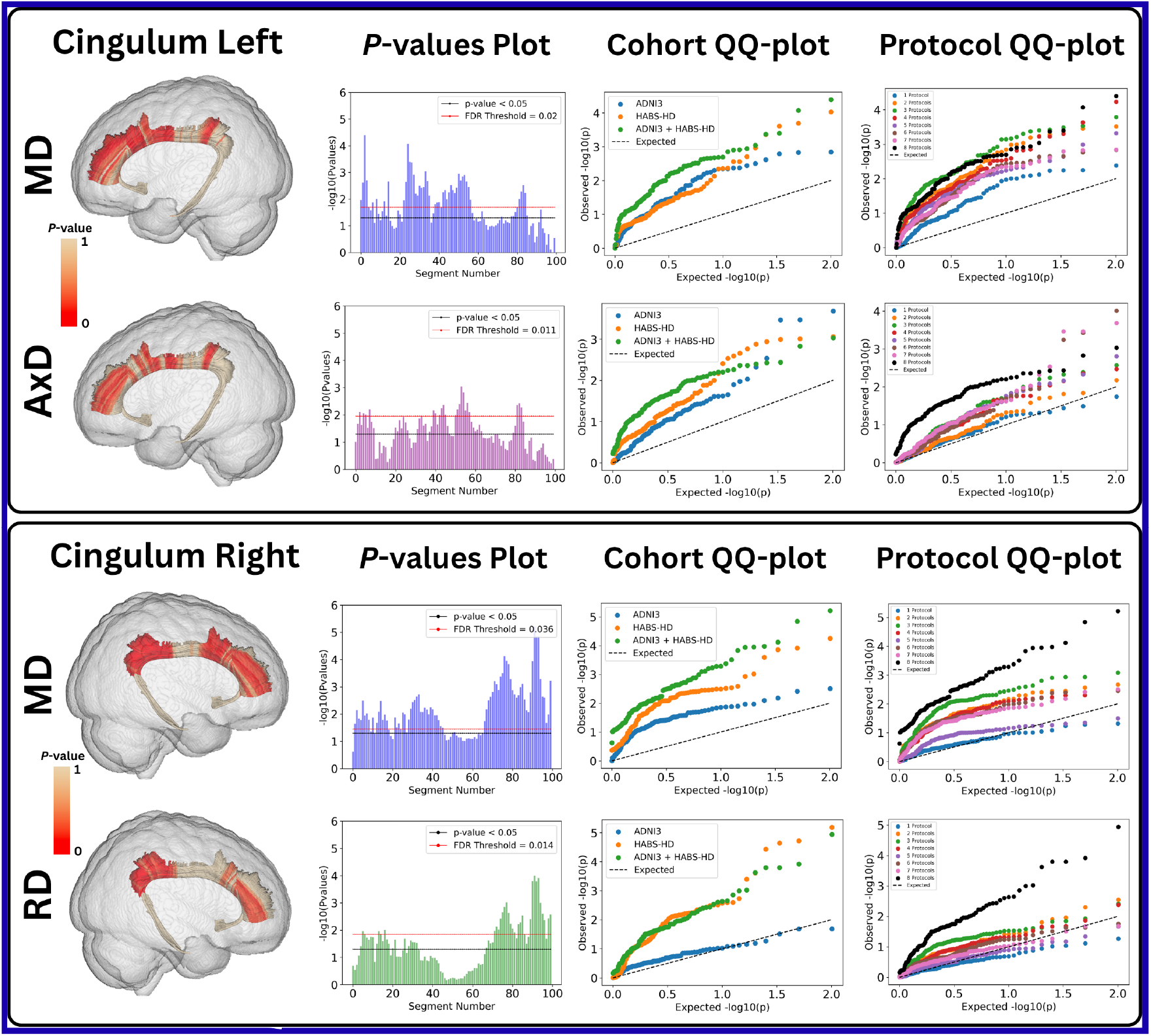
BUAN tractometry results for the cingulum bundle in the left (top) and right (bottom) hemispheres. Showing the effects of amyloid positivity. In the left cingulum, stronger effects were detected using mean and axial diffusivity (MD, AxD), while in the right cingulum, the effects were more prominent with MD and radial diffusivity (RD). The 3D tract visualizations in the first column of panels highlight regions with significant amyloid-related effects in red. *P*-value plots display the negative logarithm of *P*-value across 100 segments along each tract. Cohort QQ-plots illustrate increased signal strength when combining data from ADNI3 and HABS-HD, while protocol QQ-plots show enhanced detection power with the inclusion of multiple acquisition protocols.

## IV. Discussion

In this study, we used BUAN tractometry to examine the microstructural effects of amyloid positivity on 38 major WM tracts across participants from the ADNI3 and HABS-HD cohorts. Our results revealed widespread WM alterations associated with amyloid pathology, highlighting the potential of diffusion MRI as a non-invasive tool for understanding how the brain changes with AD pathology.

Significant differences in diffusivity metrics (MD, AxD, and RD) were observed in multiple association, projection, and commissural tracts, including the cingulum, corpus callosum, frontopontine, extreme capsule, and longitudinal fasciculi. These deviations from normality were more prominent in amyloid-positive individuals, consistent with known neurodegenerative processes associated with AD. Increased diffusivity and reduced FA, where observed, suggest demyelination and axonal degeneration, in line with prior histopathological and neuroimaging studies [4], [5], [10], [31], [32].

The cingulum bundles showed strong associations with amyloid positivity, reflecting their role in episodic memory and executive function that are typically compromised early in AD [33]. Similarly, disruptions in the corpus callosum and projection tracts such as the corticospinal and frontopontine pathways may reflect interhemispheric and subcortical–cortical disconnection, supporting the disconnection hypothesis of AD progression [34]. Beyond these, amyloid-positive individuals exhibited alterations in key association tracts, including the arcuate fasciculus, inferior fronto-occipital fasciculus, and middle and inferior longitudinal fasciculi, suggesting potential disruptions in language, semantic, and visual processing networks [35], [36]. The involvement of the extreme capsule further suggests impaired fronto-temporal communication, which may underlie executive and attentional deficits. Together, these tract-level alterations highlight how amyloid pathology may contribute to early disconnection across distributed cortical and subcortical networks, providing a mechanistic link between white-matter degeneration and cognitive decline in Alzheimer’s disease.

Notably, our analysis revealed tract-specific metric sensitivities. AxD changes were more frequent in long association fibers, while RD increases were notable in limbic and posterior tracts. These distinctions may reflect underlying pathological mechanisms, with AxD linked to axonal integrity and RD to demyelination, consistent with progressive white-matter degeneration observed in amyloid-related neurotoxicity.

Combining and harmonizing multi-site diffusion MRI data across multiple scanners and acquisition protocols enhances the generalizability of our findings. The consistent effects observed across both the ADNI3 and HABS-HD cohorts underscore the biological relevance of amyloid-related microstructural changes and demonstrate the robustness of BUAN as a tractometry framework for large-scale, multi-site studies. By integrating data from diverse sources and applying BUAN’s along-tract statistical modeling, we achieved fine-grained spatial sensitivity to localized white matter alterations; effects that may be missed by traditional voxel-based methods such as TBSS. Additionally, our QQ-plot analyses demonstrated that combining data across cohorts and protocols improves detection power, supporting the feasibility and value of harmonized tractometry analyses in multi-cohort neuroimaging research.

The observed white-matter alterations associated with amyloid positivity should be interpreted in the context of overlapping Alzheimer’s disease pathologies. Amyloid deposition likely co-occurs with tau-related neurodegeneration in these participants, as both pathologies are known to accumulate concurrently in the later preclinical and symptomatic stages of AD [37]–[40]. Therefore, the diffusion MRI changes attributed to amyloid may also partially reflect downstream or concomitant tau effects on white-matter integrity. Moreover, coexisting pathologies such as cerebrovascular disease, TDP-43 inclusions, and *α*-synuclein deposition are common in aging and Alzheimer’s disease and can further contribute to white-matter degeneration [41]. Future multimodal studies integrating both amyloid and tau PET, as well as vascular and post-mortem neuropathological markers, will be essential to disentangle their distinct and interactive contributions to tract-specific degeneration.

From a clinical perspective, identifying tract-specific white-matter alterations associated with amyloid pathology may help pinpoint early network disruptions that precede cognitive decline. Such localized microstructural changes could serve as potential imaging biomarkers for early detection, disease staging, and treatment monitoring. Although we did not stratify by clinical diagnosis in the present work, the BUAN framework and harmonized multi-cohort design provide a foundation for future analyses comparing *Aβ*+ and *Aβ*-subgroups within cognitively normal, MCI, and AD populations. These extensions could clarify whether amyloid-related white-matter degeneration follows distinct trajectories across disease stages, informing precision-medicine approaches and patient-specific therapeutic targeting.

## V. Conclusion

Our findings reveal specific white matter pathways that are susceptible to amyloid-related degeneration. BUAN tractometry offers a powerful approach for detecting spatially localized microstructural changes, providing valuable insights into how the brain is affected by Alzheimer’s pathology.

## Acknowledgment

This work was supported by the NIH (National Institutes of Health) under the AI4AD project grant U01 AG068057, grant numbers and RF1 AG057892, the National Institute on Aging (NIA) under grants RF1 NS136995, R01 AG087513, R01 AG054073, R01 AG058533, R01 AG070862, P41 EB015922 and U19AG078109, National Institute of Biomedical Imaging and Bioengineering under award numbers R01 EB027585 and R01 EB017230, and the US Alzheimer’s Association under grant AARG-23-1149996.

## Notes

https://adni.loni.usc.edu/data-samples/adni-data/

https://apps.unthsc.edu/itr/our

https://docs.dipy.org/dev/interfaces/buan_flow.html

### Competing Interest Statement

The authors have declared no competing interest.

### Summary of Updates

Added more information in the discussion section regarding the interpretation of results.

